# *twist1a* and *twist1b* play overlapping roles during zebrafish pectoral fin development

**DOI:** 10.64898/2026.09.17.752508

**Authors:** Deepam Gupta, Abby C. Lee, Deborah Yelon

**Affiliations:** Department of Cell and Developmental Biology, School of Biological Sciences, University of California, San Diego, La Jolla, CA, 92093, USA

**Keywords:** forelimb, limb bud, Twist1, Tbx5, Fgf10, Saethre-Chotzen Syndrome

## Abstract

**Background:** In humans, haploinsufficiency of *TWIST1* leads to Saethre-Chotzen Syndrome, characterized by limb abnormalities and craniofacial defects. In mouse, *Twist1* promotes limb bud outgrowth via maintenance of the apical ectodermal ridge (AER), in addition to its craniofacial roles. In zebrafish, two *twist1* paralogs, *twist1a* and *twist1b*, are known to influence cranial neural crest and cranial suture development but have not yet been implicated in limb formation.

**Results:** We find that *twist1a* and *twist1b* act redundantly during zebrafish pectoral fin development. While *twist1b* mutants have normal fins, *twist1a* mutants have small fins, and *twist1a;twist1b* double mutants lack fins. These defects appear to originate with impaired establishment of the pectoral fin field, followed by failure of AER formation, likely due to reduced *fgf10a* expression in the fin mesenchyme. Anterior-posterior patterning of the fin bud is relatively normal in *twist1a* mutants and *twist1a;twist1b* double mutants, albeit with an anterior extension of *shha* expression in some embryos.

**Conclusions:** Together, our results show that *twist1a* and *twist1b* play crucial and overlapping roles in zebrafish pectoral fin development, echoing roles played by *Twist1* in the mouse limb bud and highlighting an early impact of *twist1* genes on the initial formation of the zebrafish forelimb field.

## INTRODUCTION

The basic helix-loop-helix (bHLH) transcription factor Twist1 plays crucial roles in vertebrate limb development, regulating both the proximal-distal outgrowth and the anterior-posterior patterning of the limb. The consequences of reduced *Twist1* gene dosage illustrate its importance across species. In humans, *TWIST1* haploinsufficiency causes Saethre-Chotzen Syndrome, an autosomal dominant disorder characterized by digit patterning defects, which can include brachydactyly, broad great toes, and cutaneous syndactyly, along with craniofacial abnormalities.^1–5^ Mouse models exhibit digit patterning phenotypes reminiscent of Saethre-Chotzen Syndrome: specifically, mice heterozygous for a null mutation in *Twist1* display partially penetrant preaxial polydactyly in the forelimb and hindlimb postnatally.^1,3,6,7^ By contrast, mice homozygous for a null mutation in *Twist1* display a far more severe phenotype: these *Twist1* homozygotes have severely stunted forelimb outgrowth and die by embryonic day 11.5 (E11.5) due to defects in neural tube closure.^8^

The influence of *Twist1* on limb growth and digit formation originates during early steps of limb formation. In *Twist1* null mice, the forelimb bud forms, but fails to grow, and a morphologically distinct apical ectodermal ridge (AER), which is essential for normal proximal-distal outgrowth of the limb, does not appear.^8–10^ This is likely due to the lack of *Fgf10* expression in the limb mesenchyme, leading to inadequate crosstalk with the overlying ectoderm.^9,10^ Beyond its role in regulating proximal-distal outgrowth, *Twist1* also regulates the anterior-posterior patterning of the limb^9–13^, including the appropriate expression of *Shh* and *Hand2* in a posterior portion of the limb bud mesenchyme.^14–17^ *Twist1* null mice initiate reduced levels of *Shh* expression in the posterior limb bud mesenchyme and fail to maintain *Shh* expression subsequently.^9,10^ This phenotype has been attributed to the failure of *Twist1* null mice to form an AER, as signals from the AER are necessary for maintenance of *Shh* expression.^18,19^ Conditional inactivation of *Twist1* in the limb mesenchyme after limb bud formation can bypass the early requirement for *Twist1* during AER establishment; this temporally controlled loss of *Twist1* induces ectopic *Shh* and *Hand2* expression in an anterior portion of the limb bud, resulting in the formation of supernumerary digits.^11,13^ Similarly, reducing the level of *Twist1* function in hypomorphic mutants leads to an anterior expansion of *Shh* expression in the limb mesenchyme.^12^ Together, these studies indicate that precise levels of *Twist1* activity are required at specific times within the limb mesenchyme to control limb bud outgrowth and patterning.

Considering the significant roles played by *Twist1* in mouse limb development, it is surprising that roles for *Twist1* genes in teleost paired fin development have not yet been uncovered. Teleost fish and tetrapods share numerous developmental mechanisms governing appendage formation,^20^ making the zebrafish pectoral fin an excellent model for investigating the molecular regulation of vertebrate limb development. Due to genome duplication in the teleost lineage,^21–24^ zebrafish possess two *Twist1* orthologs – *twist1a* and *twist1b.*^25–27^ These genes have overlapping expression patterns in multiple tissues, including cranial mesenchyme, pharyngeal arches, and pectoral fins.^25–27^ Previous studies have used morpholinos and mutant alleles to show that zebrafish *twist1* orthologs play important overlapping roles during the development of cranial neural crest-derived structures.^28,29^ For example, while reduction of either *twist1a* or *twist1b* function led to only a subtle decrease in the amount of skeletogenic ectomesenchyme, simultaneous reduction of both *twist1a* and *twist1b* function caused severe defects in ectomesenchyme specification and substantial reductions in the formation of facial cartilage.^28,29^ However, these studies of the effects of *twist1a* and *twist1b* loss-of-function did not report any defects in pectoral fin development. It is unclear whether this is because pectoral fin phenotypes were not present, relatively subtle, or simply unexamined.

Here, we combine a pair of presumed null mutations – the newly generated promoterless *twist1a^sd65^* allele and the previously characterized full locus deletion *twist1b^bns353^* allele^30^ – to examine the roles of *twist1* genes during zebrafish pectoral fin development. Our results reveal a novel pectoral fin phenotype not previously reported in *twist1a;twist1b* double mutants and outline the overlapping roles that *twist1a* and *twist1b* play during fin bud formation. While *twist1b* mutants appear to have normal fins, *twist1a* mutants have only small pectoral fins, and *twist1a;twist1b* double mutants fail to form pectoral fins. Furthermore, shared functions of *twist1a* and *twist1b* are evident during early steps of fin bud development, including the establishment of the pectoral fin field, the formation of the AER, and the regulation of anterior-posterior patterning. Together, these roles of *twist1* genes in zebrafish highlight the conserved importance of *Twist1* during vertebrate limb development.

## RESULTS

### *twist1* genes play overlapping roles in development of multiple embryonic tissues

To examine the roles of *twist1* genes during pectoral fin development, we sought to utilize strong loss-of-function mutations, ideally null alleles, for both *twist1a* and *twist1b,* each of which is expressed in the pectoral fin bud (Fig. 1B,C).^25,31^ Because transcripts that contain nonsense mutations have the potential to trigger transcriptional adaptation and mask loss-of-function phenotypes,^32–34^ we chose to employ a pair of “transcriptless” mutant alleles. For *twist1b*, we selected the full locus deletion allele *twist1b^bns353^*, which has been previously characterized.^30^ For *twist1a*, we used CRISPR/Cas9-mediated genome editing^35^ to generate a novel deletion allele. This allele (*twist1a^sd65^*) removes 1011 bp upstream of the transcription start site, the entire 5’ UTR, and a large portion of the coding sequence, including most of the bHLH domain (Fig. 1A; see Experimental Procedures for more detail). We presume that *twist1a^sd65^*behaves as a null allele, since homozygous mutant embryos do not exhibit any detectable *twist1a* expression (Fig. 1D). We therefore combined *twist1b^bns353^* and *twist1a^sd65^* for our evaluation of the functions of *twist1a* and *twist1b*.

**Figure 1.**
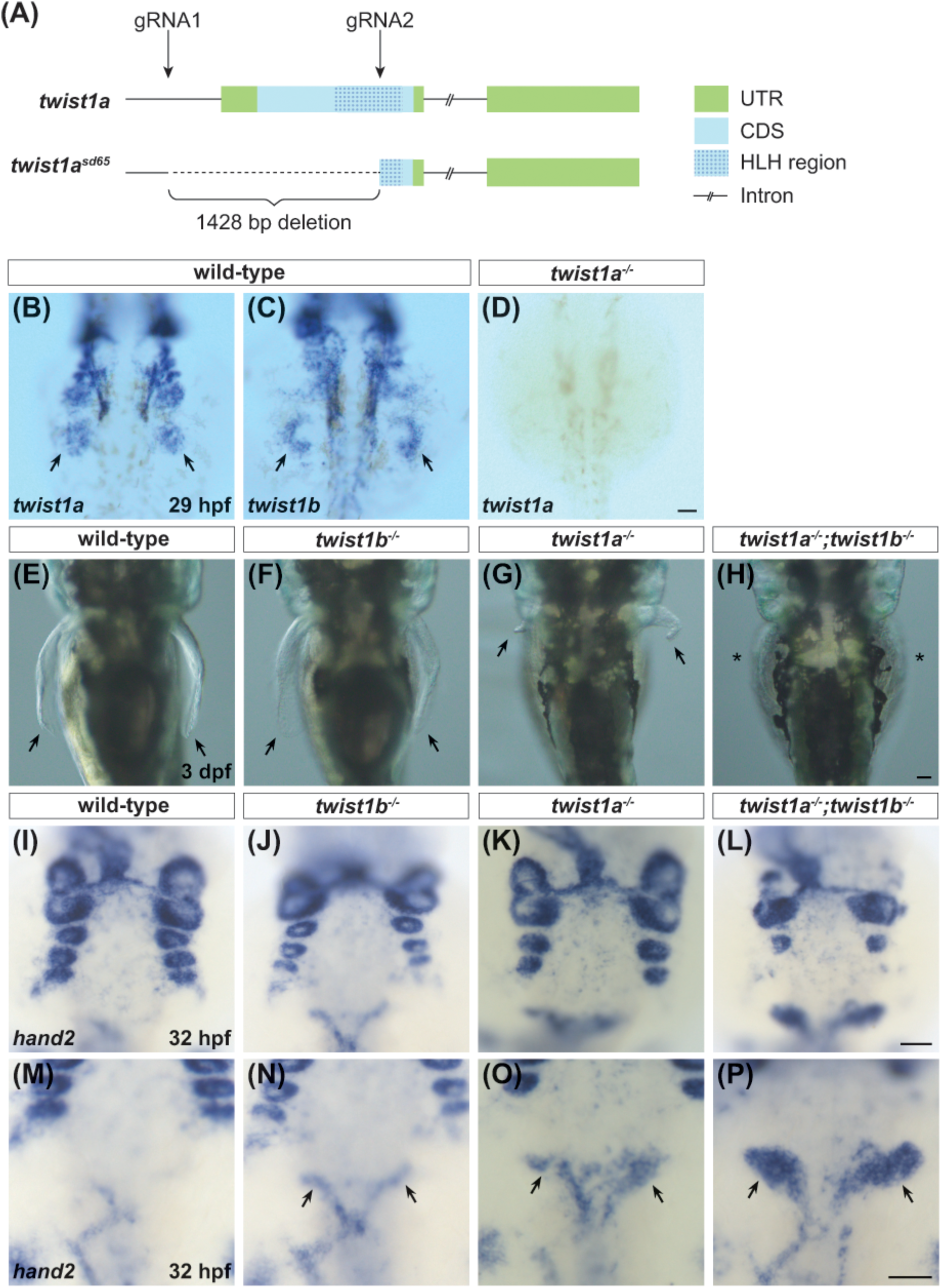
*twist1* genes play overlapping roles in multiple tissues. (**A**) Schematic of genomic location of the *twist1a* gene. Arrows point to the targets of the guide RNAs used to generate the *twist1a^sd65^* allele, a 1428 bp deletion that eliminates 1011 bp upstream of the 5’ UTR, the entire 5’ UTR, and the coding sequence for 101 amino acids. (**B-D**) Dorsal views, anterior to the top, at 29 hours post-fertilization (hpf) compare expression of *twist1a* (B) and *twist1b* (C) in wild-type embryos with expression of *twist1a* in *twist1a^sd65^* mutant embryos (D). (B,C) In wild-type, *twist1a* is expressed broadly in the pectoral fin buds (arrows, B), while *twist1b* expression is found in a subset of cells in the fin buds (arrows, C). (D) *twist1a^sd65^* mutants lack *twist1a* expression (n=6/6). (**E-H**) Dorsal views at 3 days post-fertilization (dpf) highlight the presence (arrows) or absence (asterisks) of pectoral fins. Compared to the normal pectoral fins in wild-type (E; n=6/6) and *twist1b* mutant (F; n=14/14) embryos, pectoral fins are abnormally small in *twist1a* mutants (G; n=7/7) and absent in *twist1a;twist1b* double mutants (H; n=5/5). (**I-P)** Dorsal views at 32 hpf show expression of *hand2* in the ventral portion of the pharyngeal arches (I-L) and in enteric neural crest cells (ENCCs) (M-P, arrows). Wild-type (I; n=5/5) and *twist1b* mutant (J; n=5/6) embryos exhibit five sets of clearly defined pharyngeal arches, whereas *twist1a* mutants (K; n=6/8) exhibit only four sets and *twist1a;twist1b* double mutants (L; n=7/7) have only two abnormally shaped sets. Also, while wild-type embryos (M; n=5/5) display relatively faint expression of *hand2* in ENCCs at this stage, *hand2* expression is more prominent and appears to be in more ENCCs (arrows) in *twist1b* mutants (N; n=7/8), *twist1a* mutants (O; n=7/8), and *twist1a;twist1b* double mutants (P; n=7/7). Scale bars represent 100 μm.

At 3 days post fertilization (dpf), while *twist1b^bns353^*homozygotes (hereafter referred to as *twist1b* mutants) displayed normal pectoral fins (Fig. 1E,F), *twist1a^sd65^* homozygotes (hereafter referred to as *twist1a* mutants) had highly stunted pectoral fins (Fig. 1G). These small fins contrast with the lack of reported pectoral fin phenotypes in previous studies of *twist1a^el571^* homozygotes.^29^ Furthermore, *twist1a^sd65^;twist1b^bns353^* double homozygotes (hereafter referred to as *twist1a;twist1b* double mutants) lacked any detectable pectoral fins (Fig. 1H). This striking absence of pectoral fins appears to be a novel phenotype, as no fin defects were reported in previous studies of *twist1a^el571^;twist1b^el570^*double mutants.^29^ Thus, our results provide the first demonstration that *twist1* genes are required for zebrafish pectoral fin development and indicate that *twist1a* and *twist1b* have partially redundant functions in this context.

Overlap between *twist1a* and *twist1b* function was also evident in other embryonic tissues. Since prior studies have implicated zebrafish *twist1* genes in the development of cranial neural crest-derived structures,^28,29^ we examined the impact of *twist1* gene function on *hand2* expression in the ventral pharyngeal arch mesenchyme.^36^ At 32 hours post-fertilization (hpf), *twist1b* mutants resembled wild-type, with five sets of clearly defined *hand2*-expressing pharyngeal arches, whereas *twist1a* mutants appeared slightly delayed, with just four sets of *hand2*-expressing arches at this stage (Fig. 1I-K). In *twist1a;twist1b* double mutants, however, pharyngeal arch formation was more severely disrupted, with only two sets of misshapen clusters of *hand2*-expressing cells (Fig. 1L). While examining *hand2* expression in the pharyngeal arches, we also noted interesting phenotypes near the anterior end of the gut tube in *twist1* mutants. At 32 hpf, wild-type embryos displayed faint *hand2* expression in enteric neural crest cells (ENCCs) near the foregut^37,38^, and *hand2* expression in this location was heightened in *twist1b* mutants, more intense in *twist1a* mutants, and particularly prominent in *twist1a;twist1b* double mutants (Fig. 1M-P). Together, our observations highlight multiple contexts in which *twist1a* and *twist1b* play overlapping roles during zebrafish development.

### *twist1* genes promote the establishment of the pectoral fin field

We wondered whether the pectoral fin phenotypes in zebrafish *twist1a* mutants and *twist1a;twist1b* double mutants might originate with defects in the initial specification of the pectoral fin-forming field. In *Twist1* null mice at E10.5, the size of the forelimb bud is reduced and *Tbx5* expression levels have been reported to be either normal^10^ or reduced^9^. However, the role of murine *Twist1* in the initial establishment of the *Tbx5*-expressing forelimb field, before limb bud outgrowth begins, has not been investigated. To address this in zebrafish, we examined the expression of *tbx5a*, which marks the pectoral fin field and is required for fin bud formation.^39–43^

At 20 hpf, *tbx5a* was expressed bilaterally in the pectoral fin-forming region of the lateral plate mesoderm in wild-type embryos, and *twist1b* mutants exhibited a comparable expression pattern (Fig. 2A,B). In contrast, the bilateral domains of *tbx5a* expression were reduced in *twist1a* mutants and further diminished in *twist1a;twist1b* double mutants (Fig. 2C,D). Similarly, at 32 hpf, *twist1a* mutants and *twist1a;twist1b* double mutants exhibited smaller clusters of *tbx5a*-expressing fin mesenchyme, compared to the normal *tbx5a*-expressing populations in the wild-type and *twist1b* mutant fin buds (Fig. 2E-H). These data indicate overlapping roles for *twist1a* and *twist1b* in promoting the initial establishment of the pectoral fin field; these early functions of *twist1* genes likely contribute to their overall influence on pectoral fin formation.

**Figure 2.**
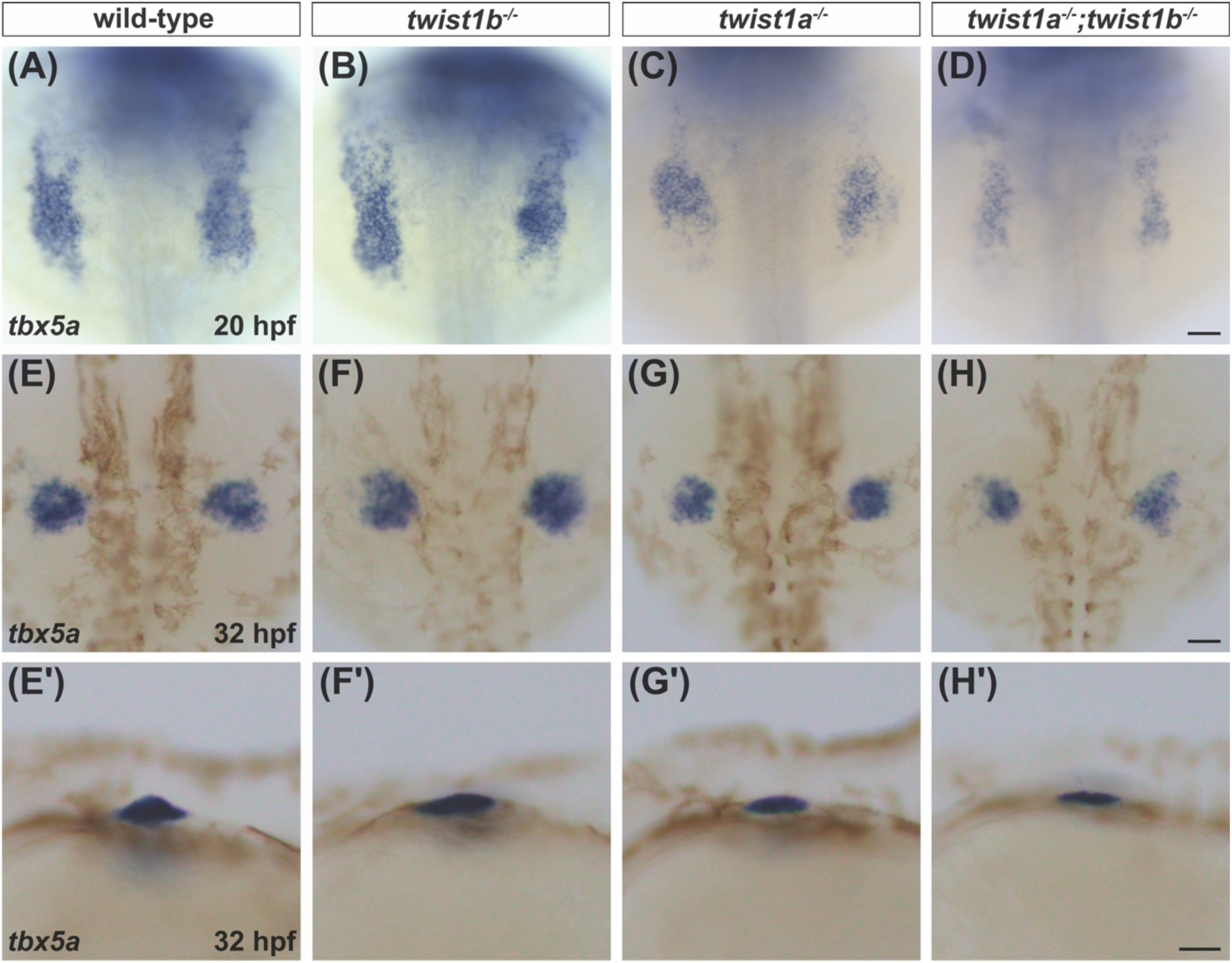
*twist1* genes promote establishment of the pectoral fin field. **(A-D)** Dorsal views show expression of *tbx5a* in the pectoral fin field at 20 hpf. This *tbx5a* expression domain is reduced in *twist1a* mutants (C; n=5/6) and further reduced in *twist1a;twist1b* double mutants (D; n=4/4), compared to the normal sizes of the domains seen in wild-type embryos (A; n=23/25) and *twist1b* mutants (B; n=19/22). **(E-H)** At 32 hpf, dorsal views, anterior to the top (E-H), and lateral views, anterior to the left (E’-H’), show that *twist1a* mutants (G; n=7/8) and *twist1a;twist1b* double mutants (H; n=5/5) continue to have smaller domains of *tbx5a* expression, compared to wild-type embryos (E; n=8/9) and *twist1b* mutants (F; n=7/8). Scale bars represent 100 μm.

### *twist1* genes play overlapping roles in AER establishment

Since the stunted forelimb outgrowth in *Twist1* null mice has been attributed to defective AER formation,^8–10^ we next examined expression of the AER marker *dlx2a*^44^ in zebrafish *twist1* mutants. At 28 hpf, expression of *dlx2a* in *twist1b* mutants resembled wild-type (Fig. 3A,B), whereas *twist1a* mutants exhibited reduced levels of *dlx2a* expression in a smaller population of cells (Fig. 3C). This reduction was further exacerbated in *twist1a;twist1b* double mutants, which displayed barely detectable levels of *dlx2a* expression (Fig. 3D). At 32 hpf, wild-type and *twist1b* mutant embryos continued to show similar patterns of *dlx2a* expression (Fig. 3E,F). In contrast, *twist1a* mutants displayed very little *dlx2a* expression (Fig. 3G), and *dlx2a* expression appeared absent in *twist1a;twist1b* double mutants (Fig. 3H). These *dlx2a* expression patterns corresponded with the morphological features of the fin bud ectoderm that were visible in live embryos. At 36 hpf, wild-type and *twist1b* mutant embryos had morphologically evident AERs of similar thickness (Fig. 3I,J), whereas the AER in *twist1a* mutants was noticeably thinner (Fig. 3K). Strikingly, *twist1a;twist1b* double mutants lacked a morphologically distinct AER (Fig. 3L). These results point to overlapping roles for *twist1a* and *twist1b* in promoting AER development. Moreover, the correlation between the severity of the AER defects and the severity of the pectoral fin morphology defects (Fig. 1G,H) in *twist1a* mutants and *twist1a;twist1b* double mutants is consistent with the well-established importance of the AER in promoting proximal-distal limb outgrowth.^45,46^

**Figure 3.**
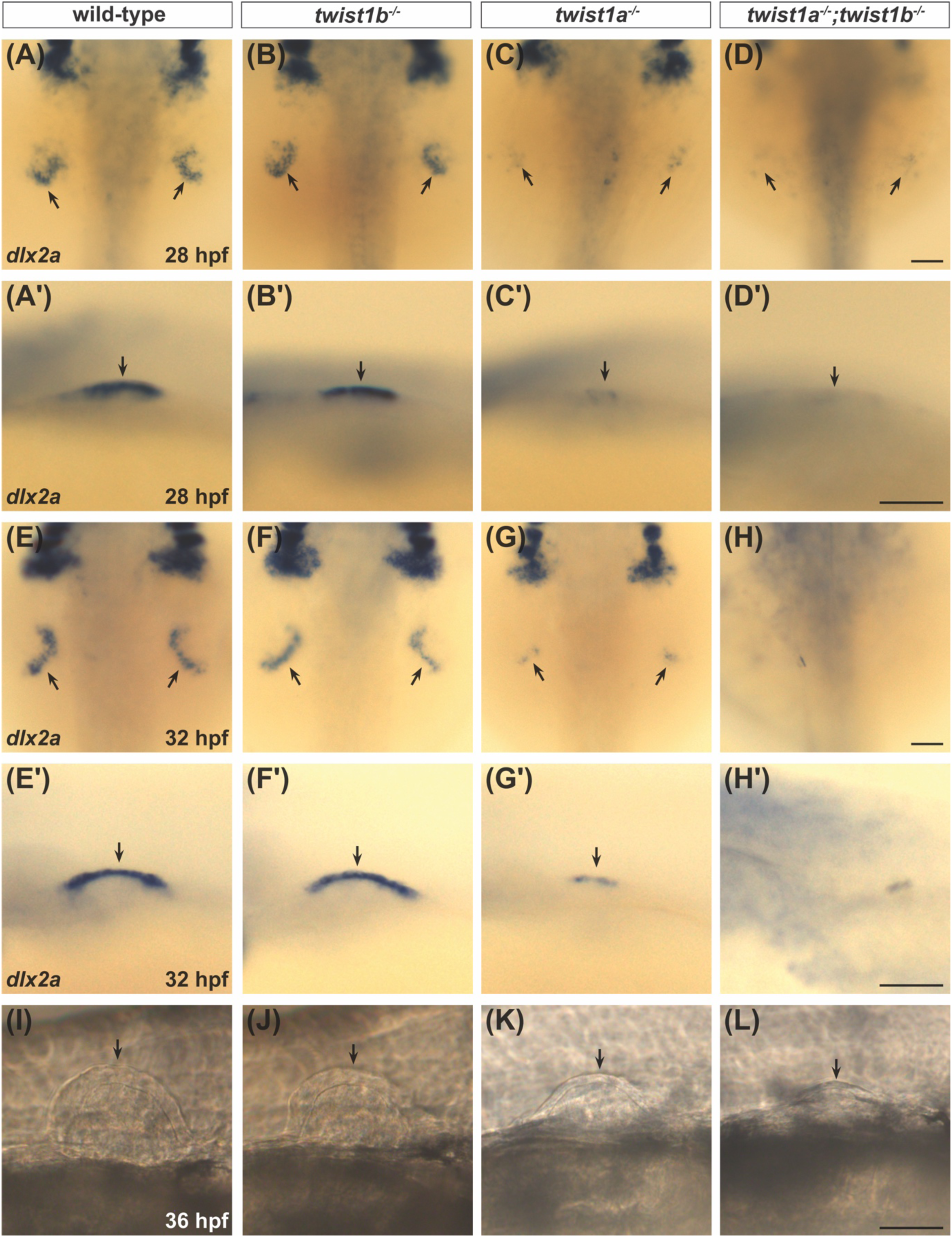
*twist1a* and *twist1b* play overlapping roles in AER establishment. **(A-H)** Dorsal (A-H) and lateral (A’-H’) views show expression of *dlx2a* in the AER (arrows) at 28 hpf (A-D) and 32 hpf (E-H). At 28 hpf, compared to wild-type embryos (A; n=12/12) and *twist1b* mutants (B; n=5/5), *dlx2a* expression is reduced in *twist1a* mutants (C; n=5/5) and further reduced in *twist1a;twist1b* double mutants (D; n=8/8). At 32 hpf, *twist1a* mutants (G; n=10/11) and *twist1a;twist1b* double mutants (H; n=7/7) continue to display reduced *dlx2a* expression, compared to wild-type embryos (E; n=9/9) and *twist1b* mutants (F; n=13/13). **(I-L)** Lateral views of pectoral fin buds in live embryos at 36 hpf show morphologically evident AERs (arrows) in wild-type embryos (I; n=5/5) and *twist1b* mutants (J; n=8/8). In contrast, *twist1a* mutants have thinner AERs (K; n=7/8), and *twist1a;twist1b* double mutants lack morphologically distinct AERs (L; n=3/3). Scale bars represent 100 μm.

To confirm that the observed defects in the *twist1a^sd65^*mutant AER are due to loss of *twist1a* function and not due to off-target effects of CRISPR/Cas9-mediated genome editing, we performed complementation testing between *twist1a^sd65^* and the previously characterized established *twist1a^el571^* mutant allele.^29^ We crossed *twist1a^sd65^* heterozygotes to *twist1a^el571^* heterozygotes and found that *twist1a^sd65/el571^*transheterozygous embryos exhibited reduced levels of *dlx2a* expression in the AER (Fig. 4D,E), compared to the levels of *dlx2a* expression in wild-type or heterozygous embryos (Fig. 4A-C). Thus, the *twist1a^sd65^* mutant allele failed to complement the *twist1a^el571^*mutant allele, consistent with both alleles disrupting *twist1a* function.

**Figure 4.**
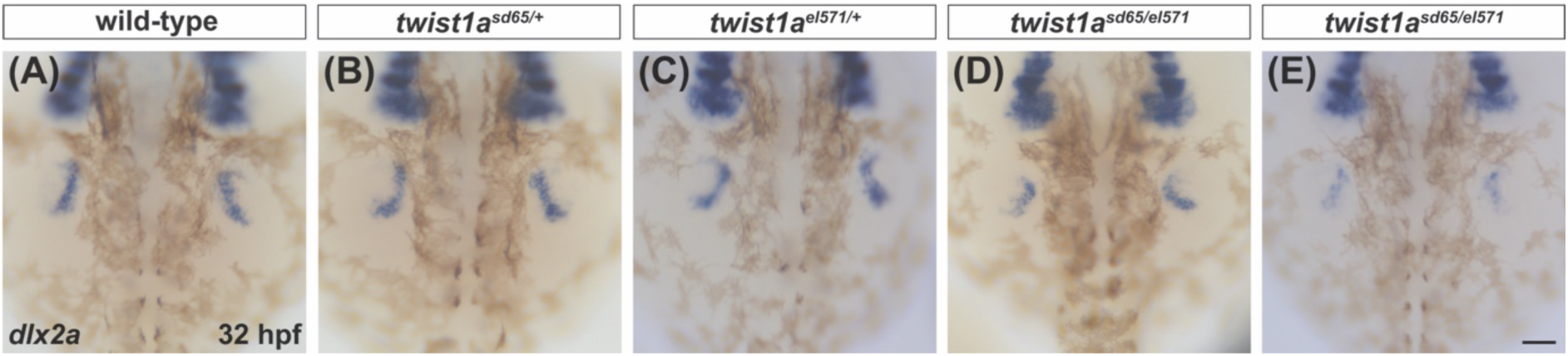
*twist1a^sd65^* fails to complement *twist1a^el571^*. **(A-E)** Dorsal views show expression of *dlx2a* at 32 hpf. Wild-type (A; n=6/6), *twist1a^sd65/+^*heterozygous (B; n=9/10), and *twist1a^el571/+^* heterozygous (C; n=14/15) embryos display similar expression patterns of *dlx2a* in the AER. However, *twist1a^sd65/el571^* transheterozygous embryos (D,E) exhibit abnormal *dlx2a* expression patterns, either resembling an earlier stage of *dlx2a* expression (D; n=10/15) similar to that shown in Fig. 3A, or having significantly reduced *dlx2a* expression (E; n=5/15). Of note, we did not observe any discernible pectoral fin outgrowth defects in transheterozygous embryos at 3 dpf (data not shown), potentially due to later recovery of delayed or reduced *dlx2a* expression. Scale bar represents 100 μm.

### *twist1* genes promote *fgf10a* expression in the pectoral fin mesenchyme

Given the AER defects found in *twist1* mutants, we investigated whether *twist1* genes promote expression of the mesenchymal factors that support AER development. In mouse, *Twist1* initiates *Fgf10* expression in the limb bud mesenchyme, which activates *Fgf8* in the ectoderm to sustain the reciprocal FGF signaling that maintains the AER.^9,10^ We therefore examined expression of *fgf10a*, which activates *fgf8a* in the zebrafish pectoral fin bud ectoderm and is required for AER development.^47^

At 30 hpf, *fgf10a* was expressed comparably in the fin mesenchyme of wild-type and *twist1b* mutant embryos (Fig. 5A,B), whereas its level and domain of expression were substantially reduced in *twist1a* mutants (Fig. 5C). The level and domain of *fgf10a* expression were further decreased in *twist1a;twist1b* double mutants (Fig. 5D). The reductions observed in *twist1a* mutants and *twist1a;twist1b* double mutants seem unlikely to result solely from a smaller amount of fin bud mesenchyme, since the contraction of their *fgf10a*-expressing domains exceeds the comparatively modest decreases in their *tbx5a*-expressing domains (Fig. 2E-H). Altogether, our results show that *twist1a* and *twist1b* play overlapping roles in promoting *fgf10a* expression, paralleling the requirement for *Twist1* in initiating *Fgf10* expression in the mouse limb bud mesenchyme and presumably underlying the role of *twist1* genes in supporting AER establishment.

**Figure 5.**
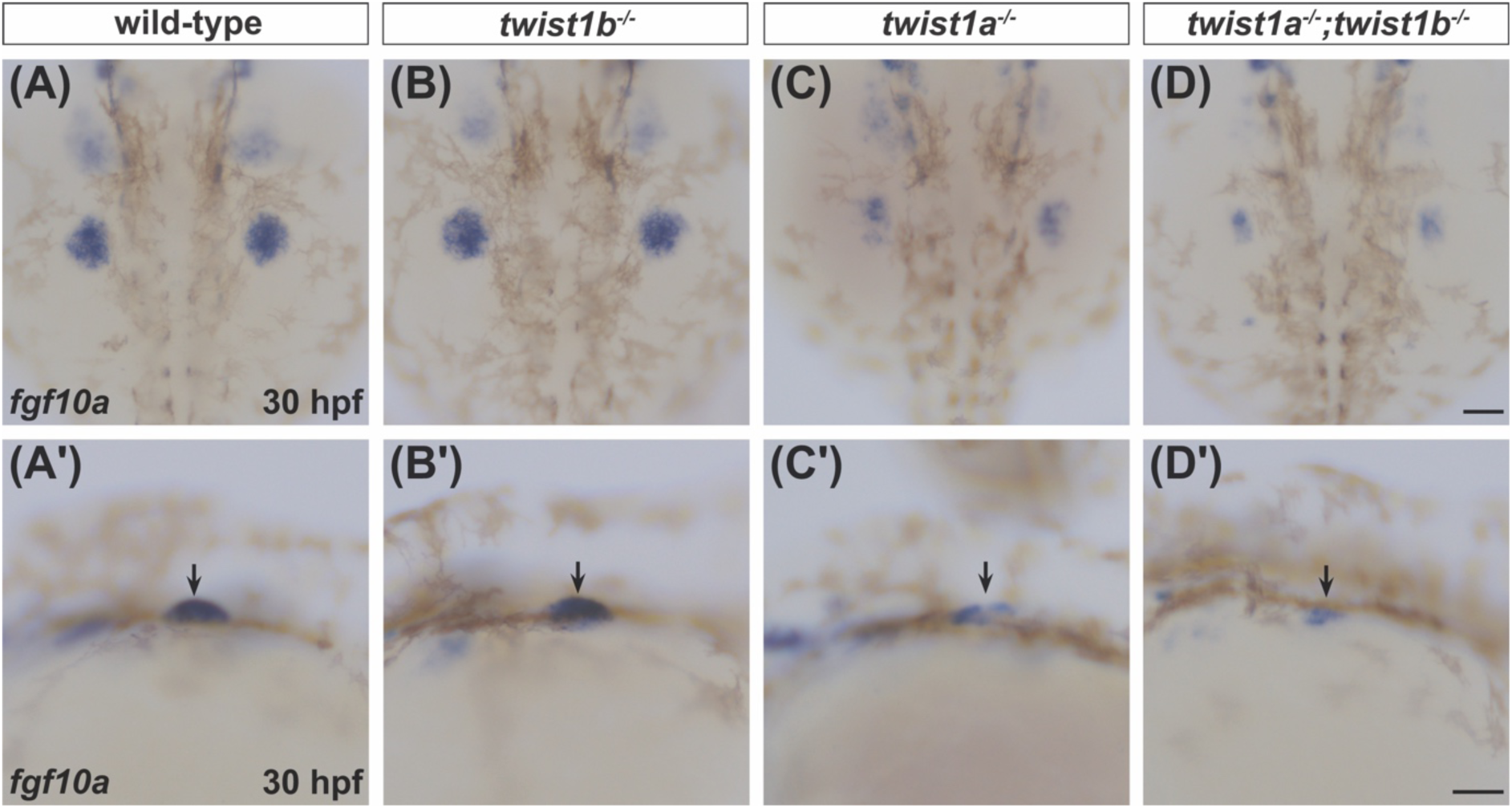
*twist1* genes promote *fgf10a* expression in the pectoral fin mesenchyme. **(A-D)** Dorsal (A-D) and lateral (A’-D’) views show expression of *fgf10a* in the pectoral fin mesenchyme (arrows, A’-D’) at 30 hpf. Compared to wild-type embryos (A; n=16/16) and *twist1b* mutants (B; n=16/16), *twist1a* mutants (C; n=11/11) and *twist1a;twist1b* double mutants (D; n=5/5) have smaller domains and reduced levels of *fgf10a* expression. Scale bars represent 100 μm.

### Anterior-posterior patterning is largely preserved, with a partially penetrant anterior expansion of *shha* expression, in *twist1* mutants

Finally, considering the role played by *Twist1* in regulating anterior-posterior patterning of the mouse limb,^9,11–13^ we investigated whether *twist1* genes contribute to the anterior-posterior patterning of the pectoral fin bud. Specifically, we examined the expression patterns of *shha* and *hand2*, both of which are normally found in the posterior fin bud mesenchyme.^48,49^

At 32 hpf, *shha* was expressed in a posterior portion of the fin mesenchyme, analogous to the tetrapod zone of polarizing activity,^48,50,51^ in wild-type, *twist1a* mutant, *twist1b* mutant, and *twist1a;twist1b* double mutant embryos (Fig. 6A-D), suggesting that *twist1* genes are not required to induce expression of *shha*. Similarly, the expression of *hand2* in a posterior portion of the fin mesenchyme appeared indistinguishable between wild-type, *twist1a* mutant, *twist1b* mutant, and *twist1a;twist1b* double mutant embryos (Fig. 6E-H), providing an additional indication that anterior-posterior patterning of the fin mesenchyme is relatively normal in *twist1* mutants. However, in a subset of *twist1a* mutants (Fig. 6C; n=6/12) and *twist1a;twist1b* double mutants (Fig. 6D; n=2/3), we observed a streak of *shha* expression extending anteriorly from the posterior domain of *shha*-expressing cells. This partially penetrant anterior extension of *shha* expression seems reminiscent of the anterior expansion of *Shh* expression observed in *Twist1* hypomorphic mice^12^ but distinct from the discrete domain of ectopic anterior *Shh* expression observed in conditional *Twist1* null mice.^11^ Together, these findings suggest that anterior-posterior patterning of the fin bud is largely preserved in zebrafish *twist1* mutants, though *twist1* genes do help to confine *shha* expression within the posterior fin mesenchyme.

**Figure 6.**
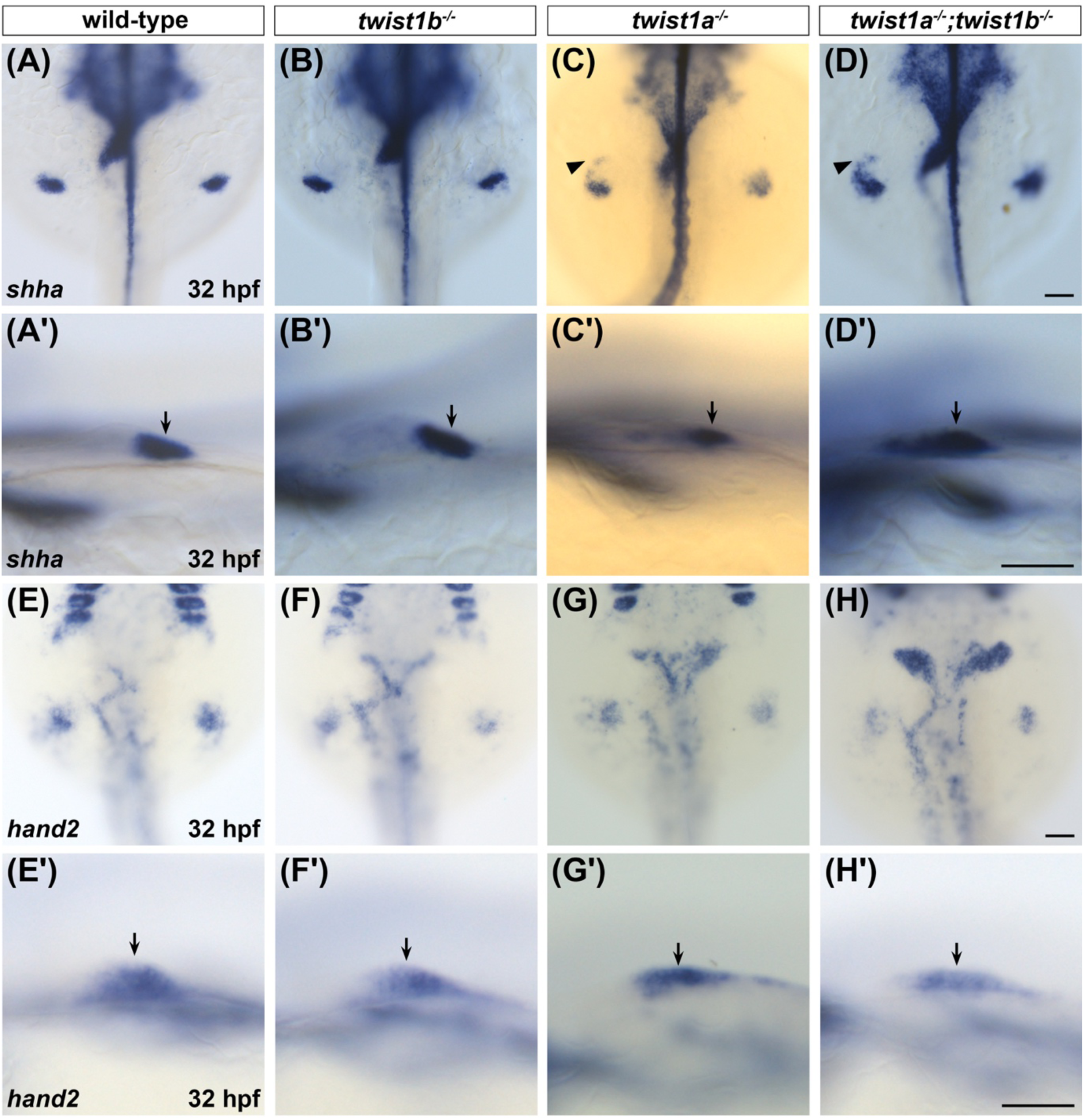
*twist1* mutants display partially penetrant anterior extension of *shha* expression and normal *hand2* expression in a posterior portion of the pectoral fin mesenchyme. **(A-D)** Dorsal (A-D) and lateral (A’-D’) views show expression of *shha* at 32 hpf. *shha* is expressed in a posterior portion of the pectoral fin mesenchyme (arrows, A’-D’) in wild-type embryos (A; n=5/5), *twist1b* mutants (B; n=6/6), *twist1a* mutants (C; n=12/12), and *twist1a;twist1b* double mutants (D; n=3/3). Whereas wild-type and *twist1b* embryos consistently display a compact posterior domain of *shha* expression, a subset of *twist1a* mutants (C, n=6/12) and *twist1a;twist1b* double mutants (D, n=2/3) also exhibit an extension of *shha* expression, reaching anteriorly beyond this domain (arrowheads, C,D). **(E-H)** Dorsal (E-H) and lateral (E’-H’) views show expression of *hand2* at 32 hpf. Wild-type embryos (E; n=5/6), *twist1b* mutants (F; n=6/6), *twist1a* mutants (G; n=7/8), and *twist1a;twist1b* double mutants (H; n=7/7) display similar patterns of *hand2* expression in a posterior portion of the pectoral fin mesenchyme (arrows, E’-H’). Scale bars represent 100 μm.

## DISCUSSION

Taken together, our results uncover a requirement for *twist1* gene function during zebrafish pectoral fin development and define the overlapping contributions of the *twist1a* and *twist1b* paralogs in this context. While pectoral fin development appears normal in *twist1b* mutants, *twist1a* mutants form stunted pectoral fins, preceded by small fin field size, impaired AER establishment, reduced *fgf10a* expression, and partially penetrant anterior extension of *shha* expression. In *twist1a;twist1b* double mutants, each of these phenotypes appears exacerbated, culminating in the failure of pectoral fin formation. We therefore conclude that *twist1a* is important for the regulation of pectoral fin outgrowth and anterior-posterior patterning. Additionally, *twist1a* can compensate for the loss of *twist1b*, and *twist1a* and *twist1b* share some overlapping functions during pectoral fin development. This functional overlap may reflect the overlap between the *twist1a* and *twist1b* expression patterns, since the broader expression domain of *twist1a* appears to encompass the more restricted domain of *twist1b* in the fin mesenchyme. Alternatively, *twist1a* and *twist1b* may have undergone partial subfunctionalization following genome duplication in teleosts, dividing some but not all of the functions of an ancestral *twist1* gene between *twist1a* and *twist1b.*^31^

Multiple aspects of the pectoral fin phenotypes in zebrafish *twist1* mutants parallel the phenotypes caused by loss of *Twist1* in the mouse forelimb, providing evidence that the functions of *Twist1* in paired appendage development are conserved between teleosts and tetrapods. As in *Twist1* null mice,^8–10^ zebrafish *twist1* mutants show defects in forelimb growth, AER development, and *fgf10* paralog expression in the forelimb mesenchyme. Because mesenchymal FGF signaling drives AER formation,^47^ the reduced *fgf10a* expression in zebrafish *twist1* mutants provides a plausible basis for their AER and outgrowth defects, mirroring the relationship of *Twist1* and *Fgf10* in mouse.^9,10^ Interestingly, our data also reveal a previously unappreciated early role of *twist1* genes in promoting the establishment of the pectoral fin field. Although *Twist1* is expressed at appropriately early stages in the lateral mesoderm in both mouse and chick,^52–54^ no prior reports have examined whether *Twist1* regulates the initial size of the *Tbx5*-expressing forelimb field. Thus, our studies also add new information that expands our understanding of the important early roles played by *twist1* genes during limb formation.

In contrast to the conservation of the roles of *twist1* genes in regulating forelimb growth, the influence of *twist1a* and *twist1b* on the anterior-posterior patterning of the zebrafish limb bud seems distinct from the known roles of *Twist1* in mouse. For example, while *Twist1* mutant mice fail to maintain *Shh* expression in the posterior forelimb mesenchyme,^9,10^ *twist1a;twist1b* double mutants exhibit robust *shha* expression in the posterior fin bud. Since the AER is critical for sending signals that sustain *Shh* expression,^19,55,56^ we suspect that *twist1a;twist1b* double mutants retain some residual inductive properties in their fin ectoderm, even though they lack a morphologically detectable AER. Additionally, although reduced function of *Twist1* in mice can lead to anterior expansion of *Hand2* expression within the forelimb bud,^12,13^ we did not detect a comparable anterior expansion of *hand2* in zebrafish *twist1a;twist1b* double mutants, nor did we observe a discrete domain of ectopic anterior *shha* expression comparable to the ectopic *Shh* found in *Twist1*-deficient mice.^11,12^ However, the partially penetrant anterior extension of *shha* expression in *twist1a;twist1b* double mutants, similar to the anterior expansion of *Shh* observed in *Twist1* hypomorphic mice,^12^ suggests some degree of conservation of *Twist1* function in restricting *Shh* expression to a posterior portion of the limb bud. The differences observed between the roles of mouse *Twist1* and zebrafish *twist1* genes during anterior-posterior patterning of the forelimb may reflect overlapping functions of *twist1* genes and other factors in zebrafish. In particular, it is possible that *twist3*, which is not found in mammals^27,31^ but is expressed in the pectoral fin bud in zebrafish,^25,27^ acts along with *twist1* genes to govern forelimb anterior-posterior patterning.

It is interesting to note that earlier studies of zebrafish *twist1* genes did not report a pectoral fin phenotype,^28,29^ potentially due to the specific nature of the *twist1* alleles examined. Our complementation test is informative in this regard: *twist1a^sd65/el571^* transheterozygotes display a less severe reduction of *dlx2a* expression compared to the reduction seen in *twist1a^sd65^* homozygotes, suggesting that *twist1a^el571^* could be a hypomorphic allele. In contrast to the transcriptless alleles used in our study, the previously examined *twist1a^el571^* and *twist1b^el570^* alleles^28,29^ are indel mutations resulting in premature stop codons that could potentially trigger nonsense-mediated decay of mutant transcripts and lead to transcriptional adaptation.^32–34^ Going forward, the combination of the transcriptless *twist1a^sd65^* and *twist1b^bns353^* alleles should be useful for future analyses of the roles of *twist1* genes during teleost paired appendage development. Ultimately, we anticipate that such analyses will contribute to a deeper comprehension of the developmental basis for the limb defects associated with Saethre-Chotzen Syndrome.

## EXPERIMENTAL PROCEDURES

### Zebrafish

We bred adult zebrafish carrying the established alleles *twist1a^el571^*(ZDB-ALT-190304-4)^29^ and *twist1b^bns353^* (ZDB-ALT-201012-6),^30^ as well as the novel zebrafish allele *twist1a^sd65^*, in these studies. All zebrafish work followed protocols approved by the Institutional Animal Care and Use Committees at the University of California, San Diego.

### Generation of a novel allele of *twist1a*

Guide RNAs (gRNAs) flanking the transcription start site of *twist1a* were selected using CHOPCHOP.^57^ Two target sequences were chosen that had no potential off-target sites in the genome with fewer than 3 bp mismatches: gRNA1 targeted a sequence upstream of the *twist1a* 5’ UTR (5’-TGAGGAGGATTTCGGCACACTGG-3’) and gRNA2 targeted a portion of the *twist1a* coding sequence (5’-GGTGGGGATGATTTTGCGCAGGG-3’) (Fig. 1A). Target-specific Alt-R crRNA molecules and common Alt-R tracrRNA molecules were synthesized by IDT. The crRNA:tracrRNA duplexes for each of the target sites were assembled in independent reactions as described previously.^35^ These duplexes were then mixed with Cas9 protein (Alt-R *S.p.* Cas9 nuclease v.3; IDT) to create crRNA:tracrRNA:Cas9 ribonucleoprotein complexes.^35^ We then injected ∼5 µM of each complex, in a total volume of 1 nl, into individual embryos.

Following injections, F0 embryos were raised to adulthood. F0 adults were crossed to wild-type fish to screen for germline transmission of mutations in *twist1a*. Genomic DNA was extracted from F1 embryos and primers flanking the gRNA target sites were used to detect successful deletions. After screening more than 50 F0 fish, we found three founders that harbored different deletions in *twist1a*. One specific mutation was selected for further propagation: this allele (*twist1a^sd65^*) is a 1428 bp deletion on chromosome 19, between bases 2230769 and 2232197 (GRCz11 genome assembly) (Fig. 1A). This deletion removes 1011 bp upstream of the 5’ UTR, the entire 5’ UTR, and the sequence that codes for the first 101 out of 171 amino acids, containing most of the Twist1a bHLH domain.

### Genotyping

In all figures (except Figs. 1B,C and 4A), “wild-type” refers to any of the following four genotypes: (1) *twist1a^+/+^;twist1b^+/+^*; (2) *twist1a^+/-^;twist1b^+/+^*; (3) *twist1a^+/+^;twist1b^+/-^*; or (4) *twist1a^+/-^;twist1b^+/-^*. We observed no phenotypic differences between any of these genotypes. We also did not observe phenotypic differences between *twist1a^-/-^;twist1b^+/+^* embryos and *twist1a^-/-^;twist1b^+/-^* embryos; we refer to both of these genotypes as *twist1a^-/-^* embryos or *twist1a* mutant embryos. Similarly, we did not observe phenotypic differences between *twist1a^+/+^;twist1b^-/-^* embryos and *twist1a^+/-^;twist1b^-/-^* embryos, and we refer to both of these genotypes as *twist1b^-/-^* embryos or *twist1b* mutant embryos.

For genotyping of the *twist1a^sd65^* allele, we used the following primers simultaneously in a single PCR reaction:

Forward primer: 5’-GCAGAAGGAAACCTGACTCTGC-3’

Reverse primer: 5’-GGACCTGACAGAGGAAGTCAAT-3’

Middle primer: 5’-GAGGATTCCGACAGTCCCAC-3’

PCR products of 253 bp and 233 bp correspond to the wild-type *twist1a* allele and the mutant *twist1a^sd65^*allele, respectively.

Separately, for genotyping of the *twist1b^bns353^* allele, we used the following primers simultaneously in a single PCR reaction:

Forward primer: 5’-GTACGCGTGGAATTGTATTTCAC-3’

Reverse primer: 5’-AACATGCATATGCACATATTTCCAAC-3’

Middle primer: 5’-CAACAGAGGTCATGGCTGTT-3’

PCR products of 109 bp and 90 bp correspond to the wild-type *twist1b* allele and the mutant *twist1b^bns353^*allele, respectively.

Genotyping for the *twist1a^el571^*allele was performed as previously described.^29^

### Whole-mount *in situ* hybridization

Whole-mount *in situ* hybridization was performed on embryos as previously described.^58^ The following probes were used: *twist1a* (ZDB-GENE-030715), *twist1b* (ZDB-GENE-050417-357), *hand2* (ZDB-GENE-000511-1), *tbx5a* (ZDB-GENE-991124-7), *dlx2a* (ZDB-GENE-980526-212), *fgf10a* (ZDB-GENE-030715-1), and *shha* (ZDB-GENE-980526-166).

### Imaging

Images of embryos following *in situ* hybridization were captured on a Zeiss Axiozoom microscope with a Zeiss Axiocam camera. Images of live embryos were captured on either a Zeiss Axiozoom microscope (Fig. 1E-H) or a Zeiss Axioimager microscope (Fig. 3I-L) with a Zeiss Axiocam camera. After imaging, individual embryos were collected in separate tubes, and genomic DNA was extracted for genotyping.

### Replicates

All observations were made in at least two independent clutches and at least two independent experiments. The numbers of embryos analyzed for each genotype with each assay are provided in the corresponding figure legends.

## ACKNOWLEDGMENTS

We thank Didier Stainier and Gage Crump for providing zebrafish strains; Gage Crump, Rosa Uribe, and members of the Yelon laboratory for helpful discussions; and Alora Yarbrough for excellent fish care.

## AUTHOR CONTRIBUTIONS

DG, ACL, and DY designed these studies; DG and ACL performed experiments; DG, ACL, and DY analyzed the data; and DG, ACL, and DY wrote the manuscript.

